# Dual-view jointly learning improves personalized drug synergy prediction

**DOI:** 10.1101/2024.03.27.586892

**Authors:** Xueliang Li, Bihan shen, Fangyoumin Feng, Kunshi Li, Hong Li

## Abstract

**Background:** Accurate and robust estimation of the synergistic drug combination is important for precision medicine. Although some computational methods have been developed, some predictions are still unreliable especially for the cross-dataset predictions, due to the complex mechanism of drug combinations and heterogeneity of cancer samples.

**Methods:** We have proposed JointSyn that utilizes dual-view jointly learning to predict sample-specific effects of drug combination from drug and cell features. JointSyn capture the drug synergy related features from two views. One view is the embedding of drug combination on cancer cell lines, and the other view is the combination of two drugs’ embeddings on cancer cell lines. Finally, the prediction net uses the features learned from the two views to predict the drug synergy of the drug combination on the cell line. In addition, we used the fine-tuning method to improve the JointSyn’s performance on the unseen subset within a dataset or cross dataset.

**Results:** JointSyn outperforms existing state-of-the-art methods in predictive accuracy and robustness across various benchmarks. Each view of JointSyn captures drug synergy-related characteristics and make complementary contributes to the final accurate prediction of drug combination. Moreover, JointSyn with fine-tuning improves its generalization ability to predict a novel drug combination or cancer sample only using a small number of experimental measurements. We also used JointSyn to generate an estimated atlas of drug synergy for pan-cancer and explored the differential pattern among cancers.

**Conclusions:** These results demonstrate the potential of JointSyn to predict drug synergy, supporting the development of personalized combinatorial therapies. The source code is available on GitHub at https://github.com/LiHongCSBLab/JointSyn.

## Background

Traditional and modern medicine have always utilized the combinatorial drug therapies to better treat cancers[1]. Compared with monotherapies, the drug combinations may improve treatment efficacy[2,3], reduce toxicity and side effects [4,5] and decrease the drug resistance [6,7]. However, responses to the same drug combination may vary largely among patients due to the high heterogeneity of cancers[3]. How to select suitable drug combination for an individual is a key challenge in personalized cancer therapy[8]. With the development of high-throughput drug combination screening in recent years[9], the responses of drug combinations can to be tested on multiple cancer cell lines simultaneously[10]. However, exhaustive searching of the entire combination space is impossible due to high costs and time consumption for the exponential increase of potential combinations[11,12]. Therefore, computational methods are needed to discover candidate synergistic drug combinations for experimental validation.

O’Neil et al have published a large-scale drug combination study, including 22,737 experimental measurements of 38 drugs on 39 cancer cell lines[13]. NCI have published a larger study of drug combination, including 304,549 experimental measurements of 104 drugs on 60 cancer cell lines[14]. Based on these observed responses, a synergy score is calculated for each “drug1-drug2-cell line” and the combination for the cell line can be classified as synergistic (observed combination effect is higher than the expected effect), antagonistic, or additive. DrugComb collected multiple scattered experimental datasets and used the unified process to calculate synergy scores[15]. In addition, pharmacogenomic databases such as Cancer Cell Line Encyclopedia (CCLE) provided comprehensive molecular measurement of cancer cell lines, including genomic mutations and copy number variations, RNA and microRNA expression profiles[16]. These resources provide a solid data basis for developing computational models to predict sample-specific drug synergy from molecular data.

Before 2018, some machine learning methods were used to predict drug synergy, such as Bayesian Network[17], Logistic Regression[18], Random Forest[19,20] and XGBoost[21–23]. In recent years, more and more deep learning methods have been developed[8,24]. Preuer et al. built DeepSynergy based on deep neural network (DNN), which uses chemical information from the drugs and genomic information from cell lines as input information, and conical layers to model drug synergies[25]. Sun et al. constructed DTF based on the tensor factorization method, which extracts latent features from drug synergy information[26]. Jiang et al. proposed a graph convolutional network (GCN), which needs to construct cell line-specific drug-drug combination, drug-protein interaction, and protein–protein interaction networks, and uses the GCN to obtain the vector representation of nodes[27]. Zhang et al. built AuDNNsynergy which was trained using all tumor samples from The Cancer Genome Atlas[28]. Wang et al. proposed DeepDDS based on GCN, which transforms the drug to molecular graph[29]. Liu et al. proposed HypergraphSynergy, which converts the drug synergy task to the link prediction task and learns drug and cell line embeddings from hypergraphs[30]. Hu et al. proposed DTSyn based on the transformer model, which extracts cell line expression profiles information with gene embedding[31].

Although the above methods have achieved satisfactory performance under cross-validation with randomly splitting, but their accuracy decreased significantly for unseen drugs and cancer samples, especially for cross-dataset prediction[29]. With the development of various transfer learning technologies in computational fields[32,33], we supposed that fine-tuning may improve the accuracy of drug synergy prediction on new drugs and cancer samples even for a small experimental dataset. Additionally, current methods modeled two drugs independently, did not sufficiently utilize the association between two drugs. Multidrug representation learning have been shown to be efficient in predicting drug-drug interactions[34], the similar idea may also promote drug synergy prediction.

Based on these, we proposed a novel deep learning model JointSyn to predict the personalized synergistic effect of drug combination. It improves from previous models by dual-view joint representation of drugs and cell lines, also fine-tuning for better robustness. JointSyn is overall the best performer on the benchmark datasets for both regression and classification tasks, for new drugs or cell lines within a dataset or cross muti-datasets. Finally, application of JointSyn to large-scale tumor cell lines obtains an estimated atlas of synergistic drug combinations for pan-cancer. Overall, both performance and case studies have proven that JointSyn is an effective tool for predicting the drug synergy, and our study also can provide quantitative suggestions for better experimental design.

## Methods

### Data

#### Drug synergy dataset

We downloaded two large-scale drug synergy datasets (O’Neil, NCI-ALMANAC) from the DrugComb database. For each triplet (drug1-drug2-cell line), the synergy score was defined by loewe additivity (LOEWE)[15]. Some triplets had repeated measurements in the downloaded data. We deleted the triplets whose synergy scores measured in repeated experiments had inconsistent signs or higher variation (coefficient of variation > 0.5), otherwise we took the median of multiple measurements as the final synergy score of this triplet. The final O’Neil dataset consists of 38 drugs, 34 cell lines and 12033 triplets[13]; The NCI-ALMANAC dataset consists of 103 drugs, 46 cell lines and 236190 triplets[14] (**Supplementary Table 1**).

#### Drug features

In order to better represent the molecular structure and physicochemical properties of the drug, we used the molecular graph and the Morgan fingerprint as drug features. SMILES of drugs were obtained from PubChem. First, RDKit was used to convert the SMILES into a molecular graph, in which the nodes are atoms and the edges are chemical bonds[35]. A 78-dimensional atomic feature[29] was calculated for each node by DeepChem[36]. Secondly, we used RDKit to convert the SMILES into Morgan fingerprints with a radius of 6[25], represented it as a 1039-dimensional binary vector[35].

#### Cell line features

Gene expression profiles and somatic mutations of cancer cell lines were collected from the CCLE database. Transcripts per million (TPM) values were log2 transformed and normalized[37]. A previous work PaccMann reported 2128 drug sensitivity-related genes screened from expression profiles and PPI networks[38]. Considering the intersection of 2128 genes and the CCLE gene expression profiles, expression values of 2087 genes were used as input features of cell lines.

### JointSyn

#### Model architecture

We proposed a novel deep learning method named JointSyn to predict drug synergy (**Figure 1a**). The input of JointSyn is the joint graph of the drug combination, the Morgan fingerprint of the two drugs, and the expression profile of the cell lines. JointSyn consists of two views, view 1 extracts the embedding of drug combination on cell line, and view 2 contacts the combination of drug embedding on cell line. Subsequently, the Prediction Net uses the embeddings learned from the two views to predict the drug synergy of the drug combination on the cell line. More details about JointSyn are introduced below.

**Fig. 1.**
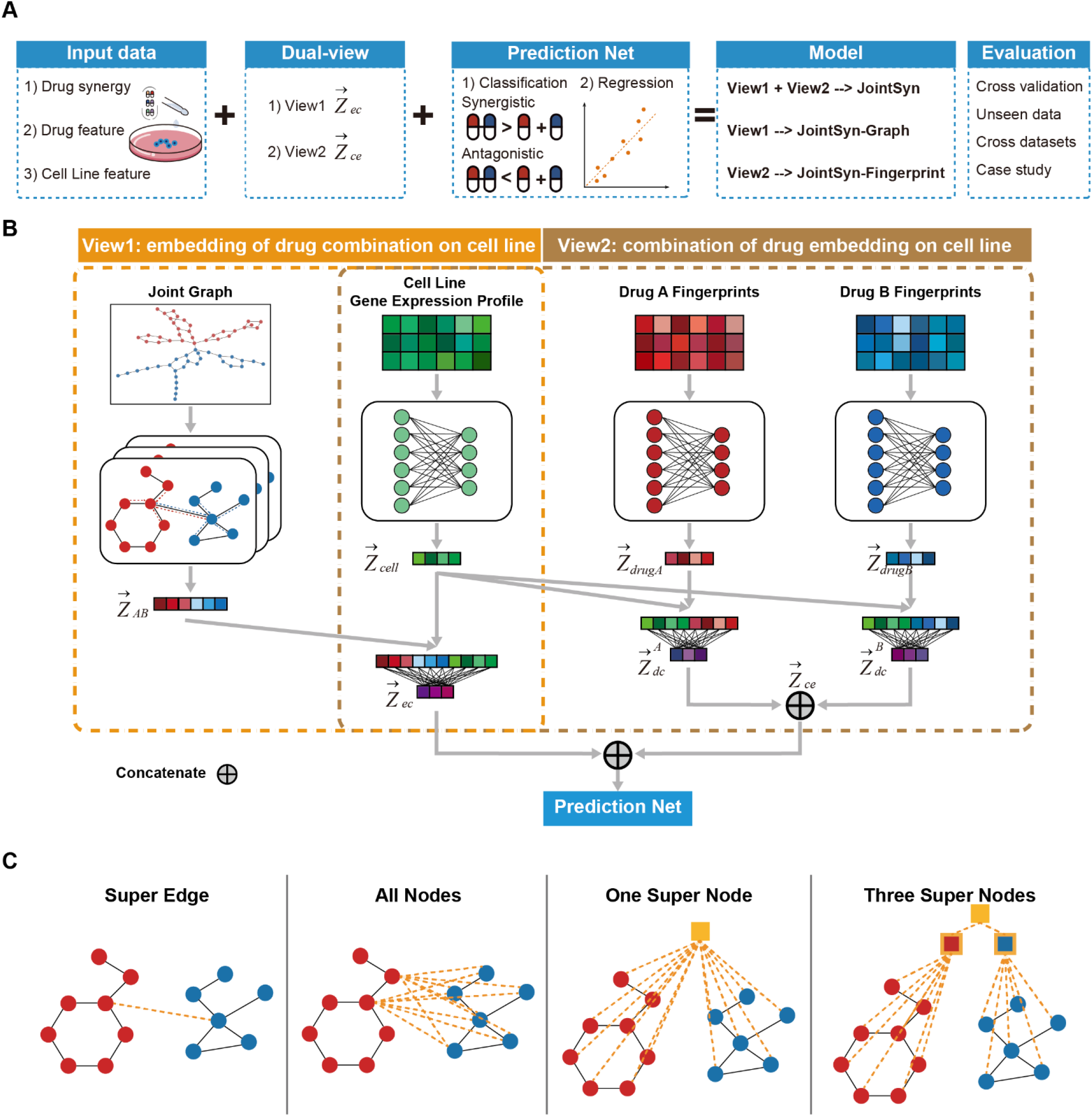
Overview of JointSyn. **a**, The schematic diagram of JointSyn. JointSyn uses drug synergy, drug features and cell line features as input, extracts drug synergy-related features from two views on classification and regression tasks. The JointSyn is evaluated by cross-validation, unseen data, cross-datasets and case study. **b**, JointSyn consists of dual views to capture the drug synergy-related features. The drug combination embedding processed through GAT based on the joint graph concatenates with the gene expression embedding to learn the embedding of drug combination on cell line. Each drug’s embedding processed through MLP based on the morgan fingerprint and concatenated with the gene expression embedding, then two drugs’ results are concatenated to learn the combination of drug embedding on cell line. **c**, Four methods for constructing the joint graph of drug combination.

#### View 1: Learning the embedding of drug combination on cell line

The molecular graph of a drug is defined as *G* = (*V, E*), where *V* is a set of *N* nodes represented by vectors, and the *i*-th atom can be represented as *ν*_*i*_ ∈ *V*; *E* is a set of edges. The chemical bond between the *i*-th and *j*-th atoms can be expressed as *e*_*i,j*_ ∈ *E*, or it can be expressed as < *ν*_*i*_, *ν*_*j*_ > ∈ *E*.

In this view, we want to get the embedding of drug combination on cell line, so the first task is to get a joint graph representation of the drug combination. As shown in **Figure 1b**, four methods were proposed to get the joint graph *G*_*AB*_ = (*V*_*AB*_, *E*_*AB*_) from drug A *G*_*A*_ = (*V*_*A*_, *E*_*A*_) and drug B *G*_*A*_ = (*V*_*A*_, *E*_*A*_).

##### Super-Edge

We firstly calculated betweenness centrality to measure the importance of a node in connecting other nodes in each drug molecular graph:

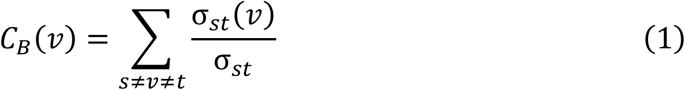

Among them, *σ*_*st*_ is the number of shortest paths from node *s* to node *t*, and *σ*_*st*_(*ν*) is the number of paths through node *ν* among these shortest paths.

Then we selected the two nodes with the highest betweenness centrality from two drug molecular graphs, namely 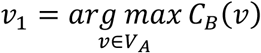 and 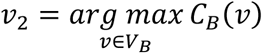, virtually added an edge between these two nodes, which was called Super Edge. Currently, *V*_*AB*_ = *V*_*A*_ ∪ *V*_*A*_, *E*_*AB*_ = *E*_*A*_ ∪ *E*_*A*_ ∪ {(*ν*_1_, *ν*_2_)}. The virtual edge can transfer atomic information between drugs and the model can capture cross-drug interactions.

##### All-Nodes

We constructed a bipartite graph 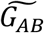 from the molecular graphs of *G*_*A*_ and *G*_*B*_, where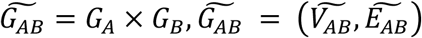. Specifically, each atom in drug *G*_*A*_ builds edges connection with each atom in drug *G*_*B*_. Currently, 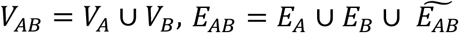.

##### One-Super-Node

We defined a super node to collect information from two drugs, and the super node is connected to each atom in the two drugs:

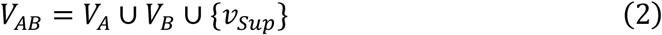

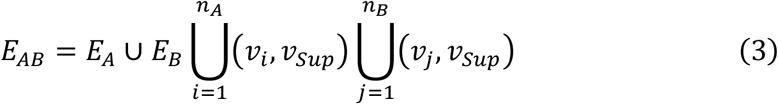

Where *n*_*A*_ and *n*_*A*_ are the numbers of atoms in drug A and B, respectively.

##### Three-Super-Nodes

We defined three super nodes to collect information about two drugs. Super node 1 (*Sup*1) is connected to all atoms in drug A, and super node 2 (*Sup* 2) is connected to all atoms in drug B. After aggregating the information of drug A and drug B, *Sup*1 and *Sup*2 are aggregated by anode *Sup*.

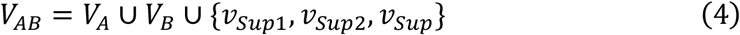

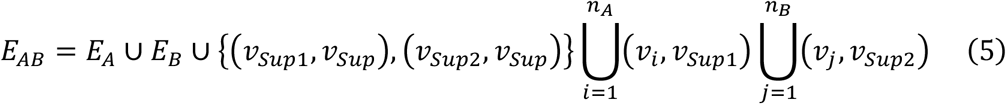

##### Extracting drug combination features based on graph attention network

We used the GAT based on the multi-head attention architecture to extract the embedding of drug combination from the joint graph *G*_*AB*_ = (*V*_*AB*_, *E*_*AB*_), whose node feature matrix is *X* and the adjacency matrix is *A*. The output features of the nodes after each layer of iterative propagation are as follows:

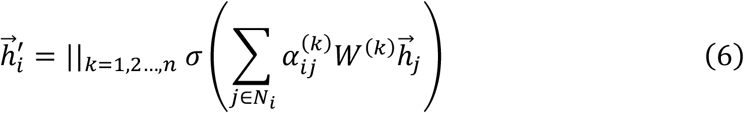

Where *k* is the number of attention heads, ∥ concatenates the outputs of multiple attention mechanisms, and *W*^(*k*)^ is the weight matrix that can be learned. The attention coefficient 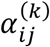 between each atom *i* and its neighbor atom *j* is calculated as follows:

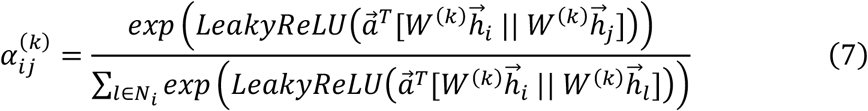

Where *W*^(*k*)^ is the weight matrix shared with the above, the purpose is to enhance the features of the vertices, || concatenates the enhanced features of vertices *i* and *j*, a 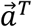 is the weight vector that can be learned, *T* is the transpose operation, and *LeakyReLU* is the nonlinear activation function.

Our model is based on a three-layer GAT, which are connected by *ReLU* activation function. After passing the three-layer GAT, each atom can see its three-hop neighbors, and through the joint graph we constructed, atomic information can be transferred between drugs. After the last layer of GAT, we add a global pooling layer to aggregate the learned multiple atomic features to obtain the embedding 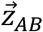 of the drug combination.

##### Extracting cell line features based on multi-layer perception

We used a two-layer MLP to construct embedding for the cell line as follows:

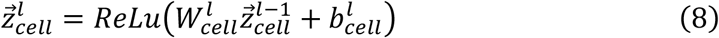

Where *W*_*cell*_ and *b*_*cell*_ are weight matrices that can be learned, *l* means in the *l*-layer. In the first layer, 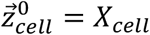 is the expression profile of 2087 genes, and *ReLU* is the nonlinear activation function. Finally, the embedding 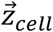 of the cell line is obtained.

##### Concatenating the embedding of drug combinations on cell lines

The drug combination embedding 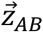 is obtained through GAT, and the cell line embedding 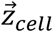 is obtained through MLP. After contacting them, the embedding of drug combination on cell line 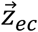 can be obtained through the fully connected layer:

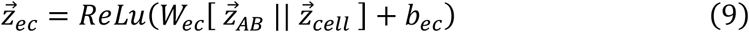

Where *W*_*ec*_ and *b*_*ec*_ are weight matrices that can be learned.

#### View 2: Learning the combination of drug embeddings on cell lines

Each drug’s embedding 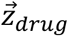 from 1039-dimensional Morgan fingerprint feature was firstly concatenated with the cell line embedding 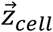, and then inputted into the MLP to obtain the embedding of one drug on the cell line 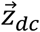:

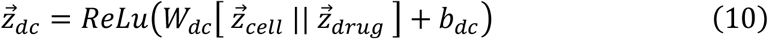

We contacted 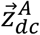 and 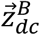 of the two drugs, and through the fully connected layer, then we can get the combination of drug embedding on cell line 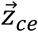:

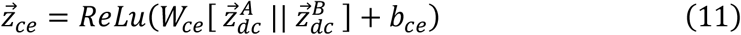

#### Prediction Net: Predicting drug synergy based on dual-view

Through the above two networks, we can get the dual-view embedding about drug synergy. By contacting these two embeddings and inputting them into the three-layer MLP:

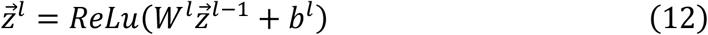

Where *W*^*l*^ and *b*^*l*^ are weight matrices that can be learned, *l* means in the *l*-layer. At the first layer,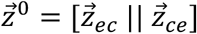, and the final embedding 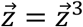.

Through the embedding 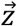 of drug synergy, we can get the final predicted value:

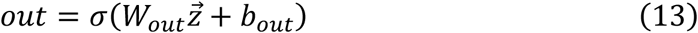

Where *W*_*out*_ and *b*_*out*_ are weight matrices that can be learned. *σ* is the softmax or linear activation function for the classification or regression task.

#### Global parameters

The architecture of JointSyn is determined by many hyperparameters, including but not limited to the loss function, activation function, learning rate and regularization method. We used grid search to adjust the hyperparameters and performed ten times of five-fold cross-validation to increase the robustness. Detailed parameter settings are in **Supplementary Table 2**.

### Method Evaluation

We compared JointSyn with several state-of-the-art methods (DeepSynergy, AuDNNsynergy, DeepDDS, DTSyn, HypergraphSynergy and XGBoost) by five-fold cross validation. Four-folds were selected as training set, and the leaved one-fold was selected as testing set which was used to evaluate the model performance. In regression tasks, we adopt Pearson Correlation Coefficient (PCC) and R square (R2) as the main evaluation metric, and we also report other widely used performance metrics, including Mean Squared Error (MSE) and Root Mean Square Error (RMSE). In the classification task, we adopt Kappa and F1 as the main evaluation indicators, and we also report other widely used classic performance indicators, including Area under the ROC Curve (ROC AUC), Precision-Recall Curve (PR AUC), Balanced Accuracy (BACC), Precision and Recall. We performed ten times five-fold cross-validations randomly and reported the mean and 95% confidence intervals of the performance metrics for each method.

To ensure the generalization ability of the model on unseen data, four data splitting strategies were used and compared: Random splitting that divided all triplets into 5 folds; PairOut splitting that randomly divided all drug combinations into 5 folds to ensure that the drug combinations in the test set do not appear in the training set; CellOut splitting that randomly divided all cell lines into 5 folds to ensure that the cell lines in the test set do not appear in the training set; DrugOut splitting that randomly divided all drugs into 5 folds to ensure that the drugs in the test set do not appear in the training set.

## Results

### Overview of JointSyn

The goal of JointSyn is to more accurate and robust estimate the personalized synergistic effects of drug combinations. Its schematic diagram is shown in **Figure 1a**. Its core features are to get the global graph information on drug combinations by joint representation, and to construct two views to capture the drug synergy related features as shown in **Figure 1b**. One view is the embedding of drug combination on cancer cell lines (view1: drugCombination-cell), and the other is the combination of two drugs’ embeddings on cancer cell lines (view2: drug1-cell & drug2-cell). In view1, molecular graphs of two drugs were firstly jointed to construct a graph of the drug combination; next graph attention network (GAT) was used to obtain a joint representation of the drug combinations; then the representation of the drug combination was concatenated the gene expression profiles of cell lines. The rationale of view1 is to better get the global graph information of drug combinations. In view2, each drug’s molecular fingerprint and cell line’ gene expression profile were integrated through multi-layer perception (MLP) to get the embedding of each drug on cell line respectively, and then the embeddings of two drugs were contacted to obtain the combination of drug embeddings on cell lines. Finally, JointSyn takes the embeddings from dual views to predict continuous or categorical synergy scores by MLP. The model only using view1 or view2 embedding and connecting MLP was named JointSyn-Graph or JointSyn-Fingerprint respectively.

We tested four joint methods for connecting drugs’ molecular graphs: Super-Edge, All-Nodes, One-Super-Node and Three-Super-Nodes (**Figure 1c**, seeing methods for details). The Super-Edge strategy achieves the best performance on the O’Neil dataset (**Supplementary Table 3**). Therefore, we selected Super-Edge as the Joint Method in the following sections.

### JointSyn improves drug synergy predictions

We first compared JointSyn with five state-of-the-art deep learning methods and one classic machine learning method on two tasks (regression and classification) using the O’Neil dataset. JointSyn achieves the best performance on all evaluation indicators for the regression task (**Figure 2a, Supplementary Table 4**). JointSyn’s R square (R2) is 0.78, which is 8.9% higher than the DeepSynergy which is the first method using deep learning for predicting drug synergy, and 6.8% higher than the DeepDDS which is the best baseline method. And JointSyn’s Pearson correlation coefficient (PCC) is 0.89, which is 5.3% higher than the DeepSynergy and 4.4% higher than the DeepDDS. When treating the drug synergy prediction as a classification task, JointSyn is still significantly better than other methods for Kappa and F1 scores, although the difference of ROC-AUC (area under the receiver operating characteristic curve) among JointSyn, XGBoost and DeepDDS is not obvious (**Figure 2b**). The similar performance improvement of JointSyn is also observed on another benchmark dataset NCI-ALMANAC (**Supplementary Figure 1 and Supplementary Table 5**). Overall, these results show the powerful ability of JointSyn to predict the drug synergy.

**Fig. 2.**
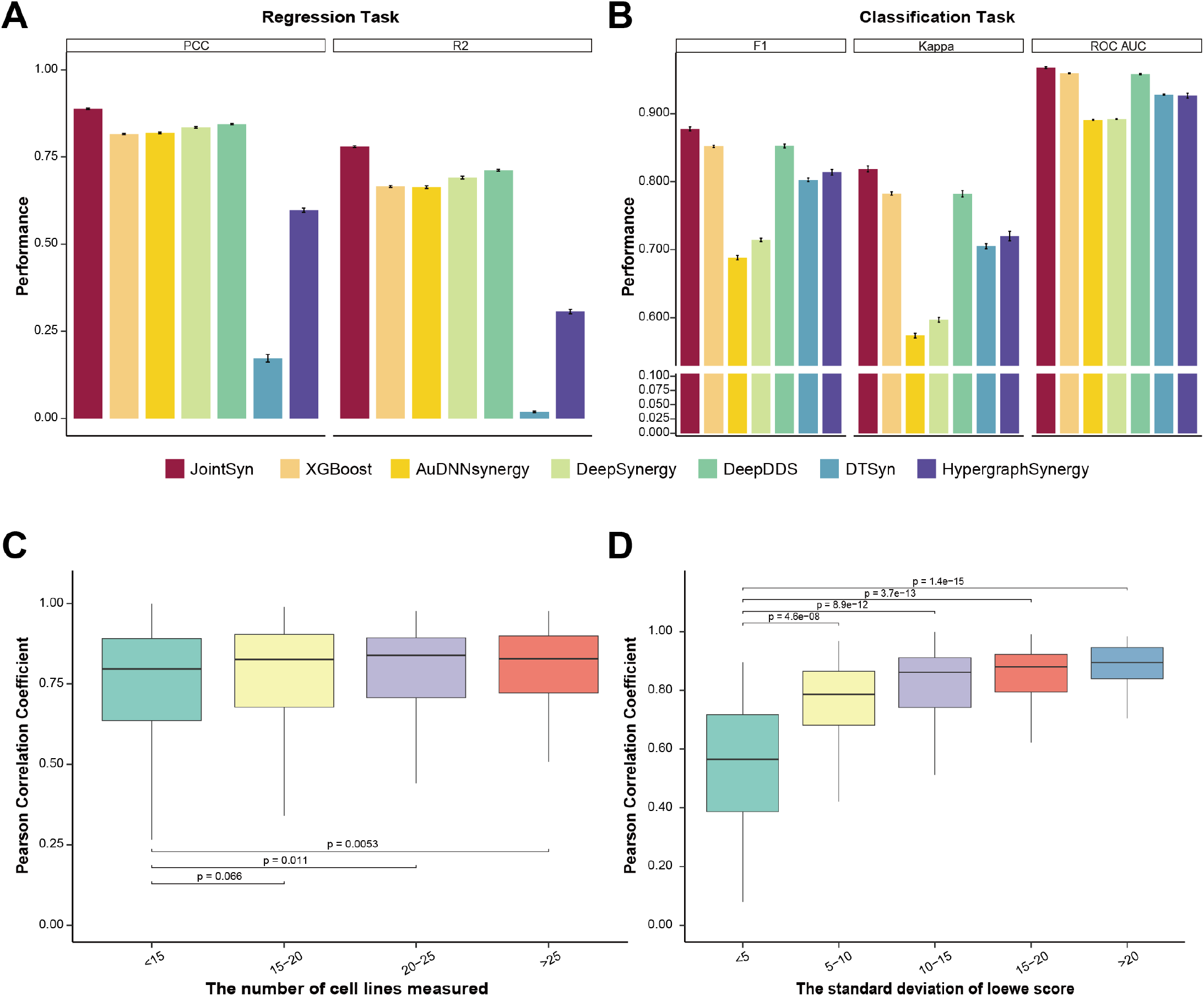
Evaluation of JointSyn using the O’Neil benchmark dataset. **a-b**, Performance comparison of JointSyn and other methods for the regression and classification tasks. Five-fold cross-validations were replicated 10 times to calculate the standard deviations (error bars). **c-d**, Taken PCC of O’Neil regression model as an example to discuss factors associated with the performance of each drug combination. **c**, The relationship between the PCC and the number of cell lines measured for each drug combination. **d**, The relationship between the PCC and the standard deviation of the real synergy scores.

There are 583 drug combinations in the O’Neil benchmark dataset, each drug combination was measured on multiple cancer cell lines. We noted that there were large differences in the performance of JointSyn for different drug combinations, therefore we further explored the possible effect factors. When a drug combination is measured in more cell lines, JointSyn’s prediction is more related with actual synergy scores (**Figure 2c**). When the standard deviation of the synergy scores for a drug combination in multiple cell lines is larger, the correlation between real and predicted synergy scores (PCC) is significantly higher (**Figure 2d**). In other words, this means that if the synergy scores of a drug combination in multiple cell lines are significantly different, the synergy scores can be predicted more accurately.

### Dual-view of JointSyn captures different aspects and achieve better performance

To inspect how well JointSyn generated the embeddings related with drug combinations on cell lines, we used t-distributed stochastic neighbor embedding (tSNE) to visualize the dual-view embeddings from the last layer of JointSyn (**Figure 3a**). Similar plots were conducted using the single view of “drugCombination-cell” embedding in the JointSyn-Graph (**Figure 3b**), and “drug1-cell & drug2-cell” embedding in the JointSyn-Fingerprint model (**Figure 3c**). The synergistic and antagonistic triples can be well distinguished in the embedding space, and the dual-view seems better than the single view. These results indicate that dual-view representation of JointSyn is successful in extract low-dimension embeddings related with drug synergy.

**Fig. 3.**
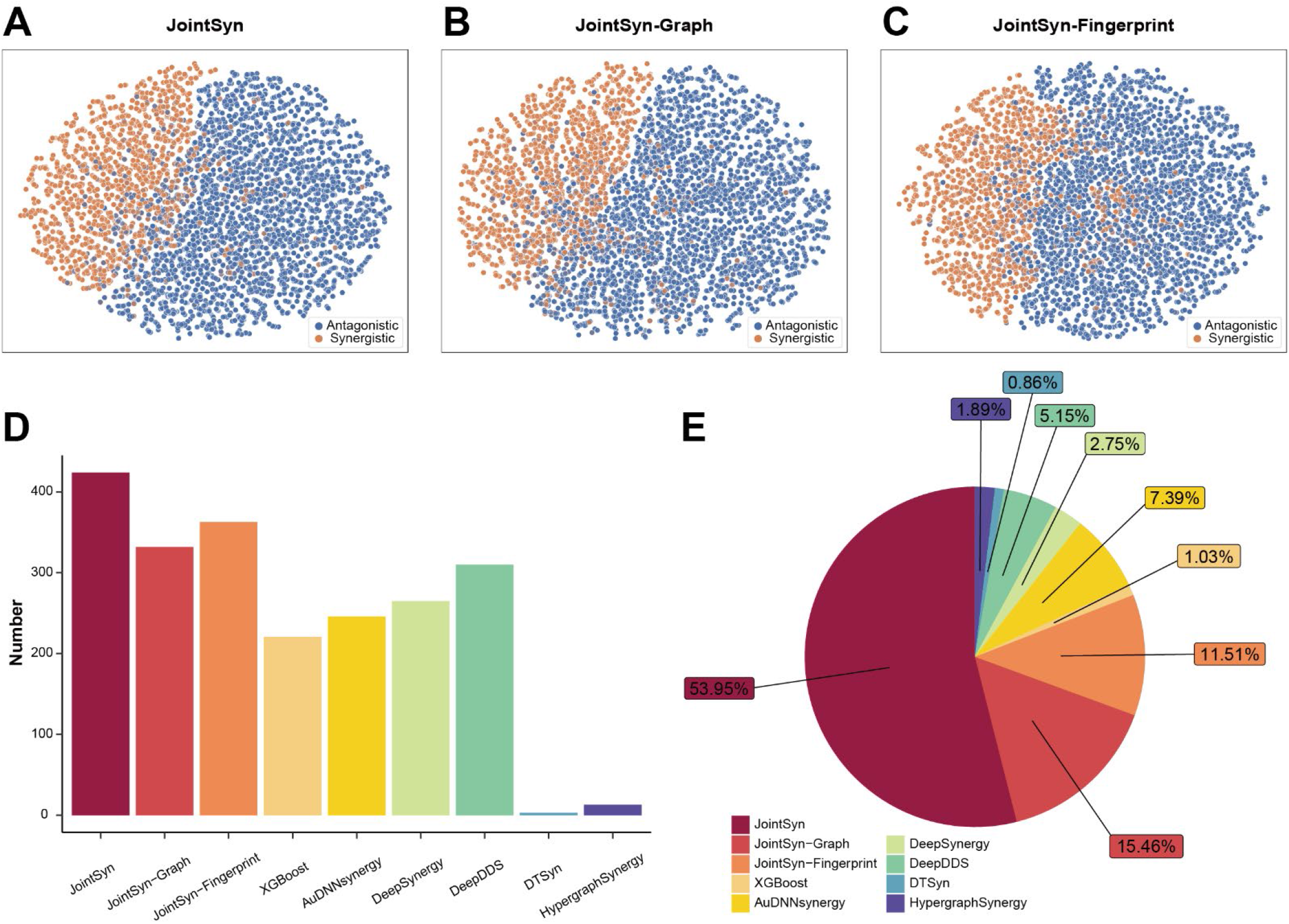
Evaluation the dual views of JointSyn. **a-c**, TSNE plots showing the last layer’ embedding from JointSyn (**a**), JointSyn-Graph (**b**) and JointSyn-Fingerprint (**c**). Each point is a triplet (drug1-drug2-cell line). Colors indicate synergistic and antagonistic triplets. **d-e**. Performance metrix was calculated for each drug combination separately instead of mix all drug combinations together. **d**, The number of drug combinations with PCC greater than 0.7 for each method. **e**, The number of drug combinations for each method achieves the best performance.

We further inspected the contribution of two views by comparing them on each drug combination. The number of drug combinations with PCC greater than 0.7 for JointSyn is 424, for JointSyn-Fingerprint is 363, and for JointSyn-Graph is 332 (**Figure 3d**). These numbers are larger than baseline methods. Next, we selected the best method (with highest PCC) for each drug combination and counted the number of drug combinations that each method obtained the best performance. The JointSyn method achieves highest PCC for 314 drug combinations, while JointSyn-Fingerprint is 67 and JointSyn-Graph is 90 (**Figure 3e**). **Supplementary Figure 2** shows the top 20 drug combinations with the highest PCC by the JointSyn, JointSyn-Graph and JointSyn-Fingerprint respectively. The best performance on some drug combinations may be achieved when using only one view. More importantly, dual-view achieves best performance for most drug combinations due to the capture of complementary aspects of embedding spaces.

### JointSyn fine-tuning improves drug synergy predictions for unseen data

In the previous part, we evaluated the performance of JointSyn by randomized data splitting, but the real prediction tasks may involve unseen drug combinations, drugs, or cell lines. Therefore, we further evaluated JointSyn by three stratified data splitting in which the drug combinations, drugs, or cell lines that were used for prediction were not included in training dataset, named PairOut, DrugOut and CellOut, respectively.

**Figure 4a** shows the performance of each method in four scenarios on the O’Neil dataset (**Supplementary Table 6** for detailed results). Compared with randomized data splitting, the performance of all methods significantly decreased for stratified data splitting. DrugOut decreases the most, followed by CellOut and PairOut. For the PairOut scenario, models may still learn from other combinations that share a drug; For CellOut scenario, models may still learn from similar cell lines; However, DrugOut simulates the prediction of a completely new drug, lack of information in the training set result in significant performance decline. Compared with other methods, JointSyn still achieves the best performance in most metrics in the stratified scenarios. For an unseen drug combination, JointSyn can averagely achieve a F1 of 0.84 and a PCC of 0.86; For an unseen cell line, JointSyn can still averagely achieve a F1 of 0.75 and a PCC of 0.67.

**Fig. 4.**
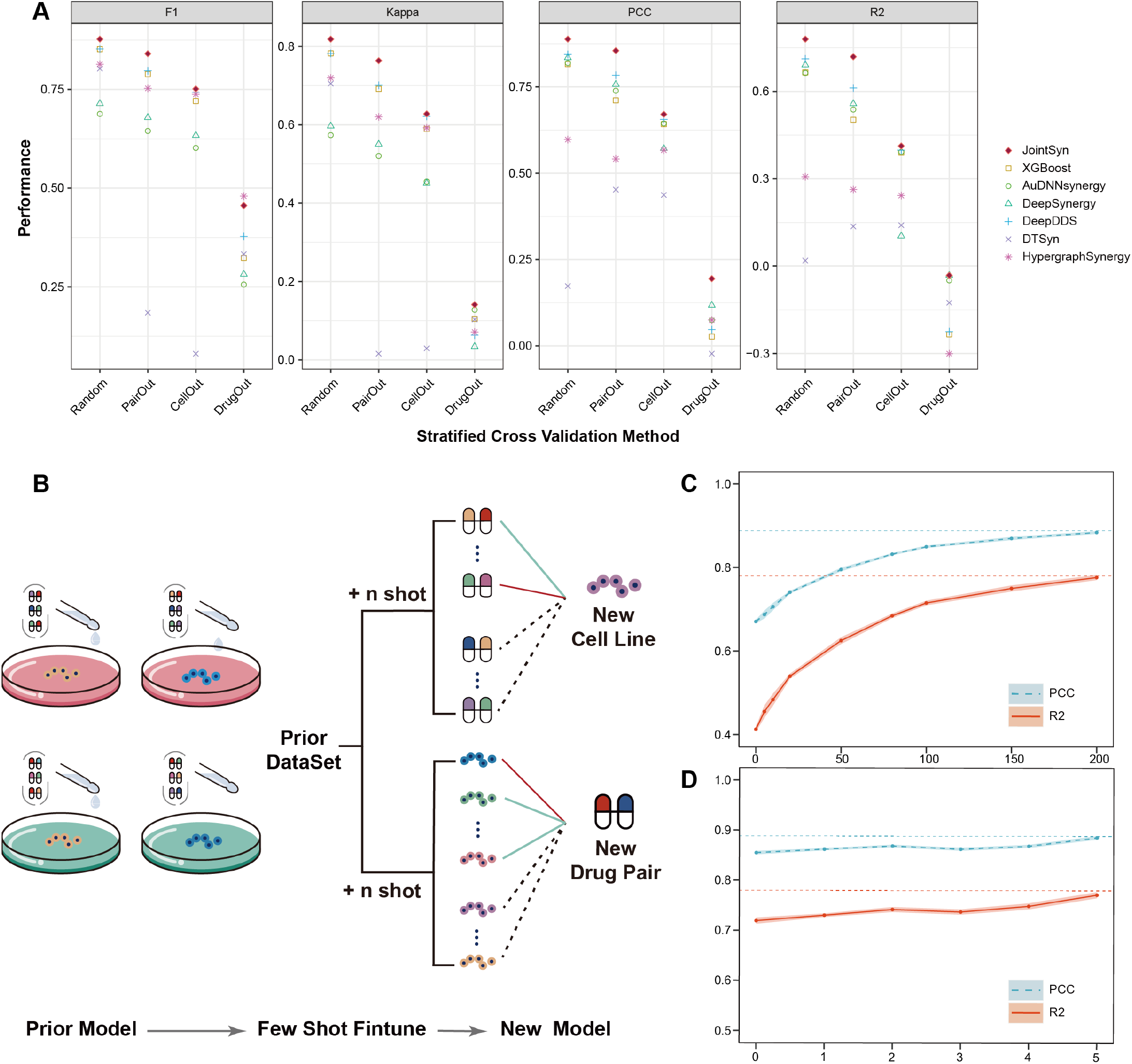
Model performance on unseen drug combinations, cell lines, and drugs. **a**, Performance comparison for the regression and classification tasks in four data splitting scenarios of the O’Neil dataset. Random means data was randomly split into training and test set. Three stratified data splitting PairOut, CellOut, DrugOut means drug combinations, drugs, or cell lines that were used for prediction were not included in training dataset. **b**, Supposing there is a new cell line (CellOut) or drug combination (PairOut), JointSyn use few-shot finetuning to improve performance. **c**, The improvement of model performance after adding experimental measurements from different numbers of drug combinations when the cell line is new. The dashed line is the JointSyn’s performance in random data splitting scenarios. **d**, The improvement of model performance after adding experimental measurements from different numbers of cell lines when the drug combination is new.

Prediction of drug synergy for unseen data is very challenging. To address this challenge, we used the fine-tuning method to improve the JointSyn’s performance by introducing a small number of experimental measurements (**Figure 4b**). For a new cell line in CellOut splitting scenario, k drug combinations on this cell line were gradually added into the training set (k-shorts). As the number of shots increased, both PCC and R2 gradually increased and tended to be stable (**Figure 4c**). The performance metrics at 150-shots is close to random splitting. This means that, for a given drug list, if 21% (150/703≈21%) combinations are experimentally measured for a cell line, and the synergy of the remaining drug combinations can be well predicted. Similarly, k-shorts fine-tuning promoted JointSyn’s performance in the PairOut scenario (**Figure 4d**). For a drug combination that is not in the training set, when the synergy scores of this combination on 15% (5/34≈15%) cell lines were added into the training set, JointSyn can well predict synergy scores on the remaining cell lines. Taken together, JointSyn with fine-tuning further improve its generalization ability for prediction with few experimental measurements.

### JointSyn fine-tuning improves drug synergy predictions for independent datasets and provides quantitative suggestions for better experimental design

We further evaluated JointSyn for cross-dataset prediction using NCI-ALMANAC and O’Neil datasets. These two datasets only shared 14 drugs, 9 cell lines and 221 triplets (drug1-drug2-cell line) (**Figure 5a**). For the overlapped triplets, whose synergy scores were measured in both datasets, the correlation of two datasets is only 0.32 (**Figure 5b**). Such low correlation may come from differences in experimental protocols between two laboratories such as the concentration range of drugs and the cell viability assay[39]. Taken the larger dataset NCI-ALMANAC (236190 triplets) as training data and the O’Neil dataset (12033 triplets) as independent testing data, the performance of JointSyn (R2 = 0.064 and PCC = 0.14) and other baseline methods significantly dropped. Such low predictive performance indicates that direct application of the model to independent datasets is infeasible due to huge differences in experimental subjects and protocols.

**Fig. 5.**
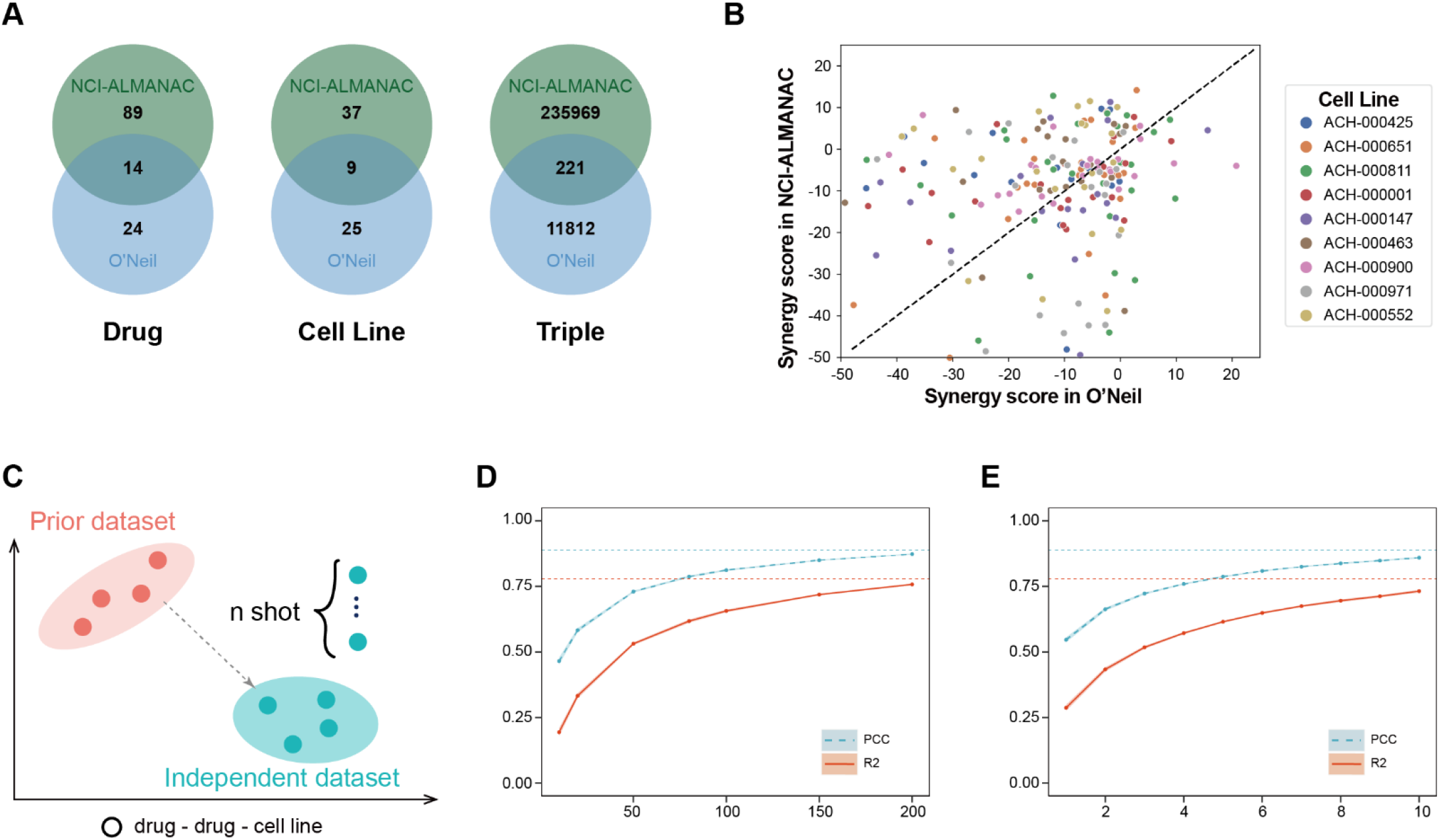
The difficulties and model performance for cross-dataset prediction. **a**, Venn diagram of drugs, cell lines and triples (drug1-drug2-cell) in O’Neil and ALMANAC datasets. **b**, A scatter plot shows the synergy scores of the overlapped measured triplets between O’Neil and NCI-ALMANAC datasets. **c**, The process of few-shot finetuning for cross-dataset. **d-e**, The performance of the model fine-tuned by the O’Neil dataset based on the model trained by NCI-ALMANAC dataset. **d**, Performance improvement with the adding of experimentally measured drug combinations on a new cell line. **e**, Performance improvement with the adding of experimentally measured cell lines on a new drug combination.

The cross-dataset prediction is a common demand in real-world application scenarios and a very challenging task for computation modelling from limited training data, but previous methods did not pay enough attention to this issue. Therefore, we quantitatively explored how much data are needed for reliable predictions when training JointSyn by a prior public dataset and fine-tuning by a small independent dataset (**Figure 5c**). **Figure 5d** shows the performance of the model fine-tuned by the O’Neil dataset based on the model trained by NCI-ALMANAC. For a given drug list, when 28% (200/703≈28 %) drug combinations for each cell line of the O’Neil dataset were added to fine-tune JointSyn, synergy scores of the remaining triplets can be well predicted (R2 = 0.75, PCC = 0.87). Similarly, when 29% (10/34≈29%) cell lines for each drug combination were added, JointSyn can well predict synergy scores on the remaining cell lines (R2 = 0.73, PCC = 0.86). This experiment proves that through fine-tuning with a small number of experimental measurements from the external dataset, the decrease of model performance on cross-dataset prediction can be solved.

### Application of JointSyn to investigate synergistic drug combinations of pan-cancers

To further demonstrate the utility of JointSyn in personalized drug synergy prediction and explore the patterns of drug synergy among different tumors, we applied JointSyn trained with the O’Neil dataset to score 996 cell lines from seven tumor lineages of CCLE on 703 drug combinations. This generated an estimated atlas of synergistic drug combinations for pan-cancer (**Figure 6a**). Ward’s hierarchical clustering divided tumors into 4 clusters (C1∼C4) and drug combinations into 5 clusters (D1∼D5) from the predicted drug synergy matrix. D1 and D2 have no synergistic effects in almost all tumors, while other drug combination clusters show heterogeneous patterns of synergy scores among different tumors. Tumors in C1 are mainly composed of blood cancers and lymphomas, harbor no synergistic drug combinations, which may reflect huge difference between hematolymphoid tumors and solid tumors; C3 is mainly composed of skin cancer; Other types of solid tumors are mixed in C2 and C4.

**Fig. 6.**
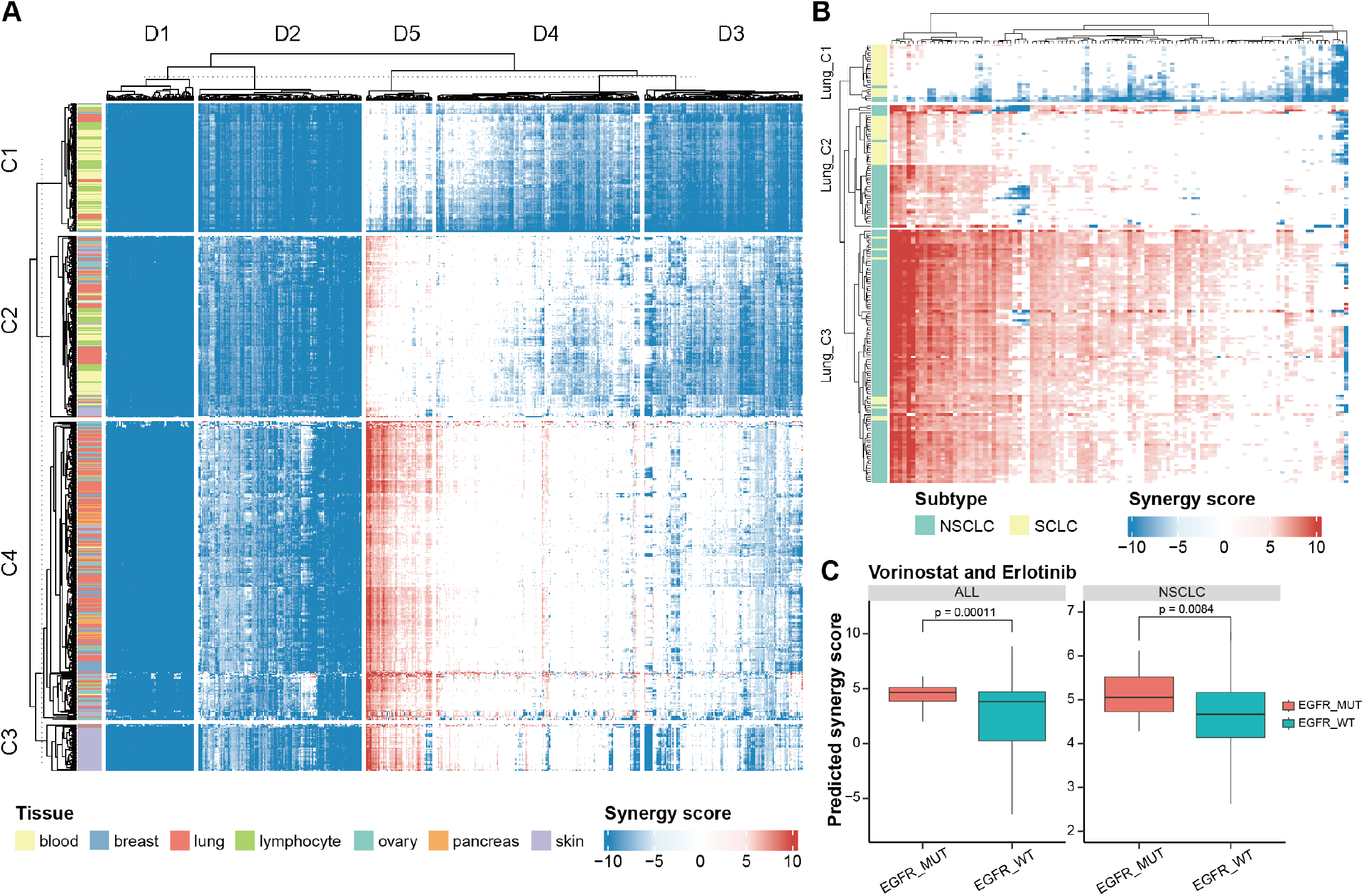
The predicted synergy scores of pan-cancers. **a**, The predicted synergy scores of 703 drug combinations on 996 cell lines. Larger positive value (red) indicates more synergism, and smaller negative value (blue) indicates more antagonism. **b**, The predicted synergy scores of selected drug combinations on lung cancer cell lines. NSCLC: Non-Small Cell Lung Cancer. SCLC: Small Cell Lung Cancer. **c**, Comparison of the predicted synergy scores of vorinostat and erlotinib in cell lines with or without EGFR mutations. Pan-cancers and NSCLC were compared respectively. P-value was calculated by the Wilcoxon test.

Lung cancer has the largest number of cell lines in CCLE dataset. Next, we taken lung cancer as an example to illustrate the heterogeneity among cancer samples of the same lineage. The drug synergy matrix for all 703 drug combinations in 188 lung cancer cell lines was illustrated in **Supplementary Figure 3a**. We selected 108 drug combinations with a synergistic ratio greater than 5% and drew a heatmap of synergy scores (**Figure 6b)**. Samples were divided into three clusters (Lung_C1∼Lung_C3) based on the synergy scores of these selected drug combinations. The two main types of lung cancer, non-small cell lung cancer (NSCLC) and small cell lung cancer (SCLC), show obvious differences in the distribution of synergy scores. Lung_C1 is mainly composed of SCLC and has almost no synergistic drug combinations. Lung_C3 is mainly composed of NSCLC and many drug combinations have synergistic effects. This is consistent with many previous studies which mentioned SCLC is highly drug-resistant and has poor prognosis[40,41].

Some drug combinations have synergistic effects on specific NSCLC cell lines, therefore we further explored whether the differences of drug synergy were related to certain somatic mutations. EGFR is the frequently mutated driver gene in NSCLC and its inhibitor erlotinib is in our training dataset. Combinations of erlotinib with some other drugs are synergistic based on the predictions from JointSyn and associated with EGFR mutations. Taken “erlotinib and vorinostat”, “erlotinib and dactolisib”, “erlotinib and MK-2206” as examples, they have significantly high synergy scores in EGFR mutated cell lines than those without EGFR mutations, whether in NSCLC cell lines or in all cell lines (**Figure 6c, Supplementary Figure 3b and Figure 3c**). Evidence related to these predicted drug synergy can be found in previous studies. Vorinostat can enhance the therapeutic potential of erlotinib in lung cancer cells[42]; the combination of MK-2206 and erlotinib can synergistically inhibit the cell proliferation of human cancer cell lines[43]; PI3K/Akt/mTOR signaling is the main mechanism of EGFR resistance, and dactolisib is a dual PI3K/mTOR inhibitor[44], which may explain the synergy effect of erlotinib and dactolisib. In summary, the predictions from JointSyn are mostly reliable and some are supported by existing experimental evidence.

## Discussion

In this work, we have proposed a novel deep learning model, JointSyn, to predict drug synergy from dual-view jointly learning. JointSyn performs best compared to other methods on two benchmark datasets. The embedding from dual view has been proved to be significantly helpful in drug synergy prediction. More importantly, JointSyn utilizes few experimental measurements to fine-tune, improve its performance not only for the unseen subset within a dataset but also for independent dataset. Finally, an estimated atlas of synergistic drug combination for pan-cancer were generated by JointSyn and the differential pattern among tumors were discussed.

A common bottleneck of developing drug synergy prediction methods is the limited number of experimental measured synergy scores. A future direction is to incorporate large-scale unsupervised pre-training into the training process, so that the model could learn more drugs and cell lines even though these are unlabeled. JointSyn currently only uses the drug’s molecular graph, Morgan fingerprint and the expression profile of cell line to predict drug synergy. More prior information such as drug-target genes, drug-drug interactions, and drug-disturbed expression profiles may also be useful. Incorporation more information may more sufficiently model drugs and cell lines, even better capture the association between drugs in the training set and novel drugs. We will improve JointSyn from these perspectives in future. We believe that JointSyn is a valuable tool for pre-screening of synergistic drug combinations.

## Conclusions

In summary, we have proposed JointSyn which utilizes dual-view jointly learning to predict drug synergy. JointSyn outperforms existing state-of-the-art methods across various benchmarks. JointSyn with fine-tuning improves its performance for the unseen subset within a dataset and independent dataset only using a small number of experimental measurements. The application of JointSyn to generate an estimated atlas of drug synergy for pan-cancer has proved the JointSyn is a useful tool to estimate the sample-specific effects of drug combination. Overall, we believe JointSyn is a valuable tool to predict drug synergy, supporting the development of personalized combinatorial therapies.

## Availability of data and materials

All source code that produced the findings of the study, including all main and supplemental figures, are available at https://github.com/LiHongCSBLab/JointSyn.

The O’Neil and NCI-ALMANAC drug synergy datasets used in this manuscript were downloaded from the DrugComb (https://drugcomb.fimm.fi/). The SMILES of drugs were downloaded from PubChem (https://pubchem.ncbi.nlm.nih.gov/). The gene expression profiles of cancer cell lines were downloaded from CCLE (https://sites.broadinstitute.org/ccle/).

