## Supplementary Figures for "Dual-view jointly learning improves personalized drug synergy prediction"

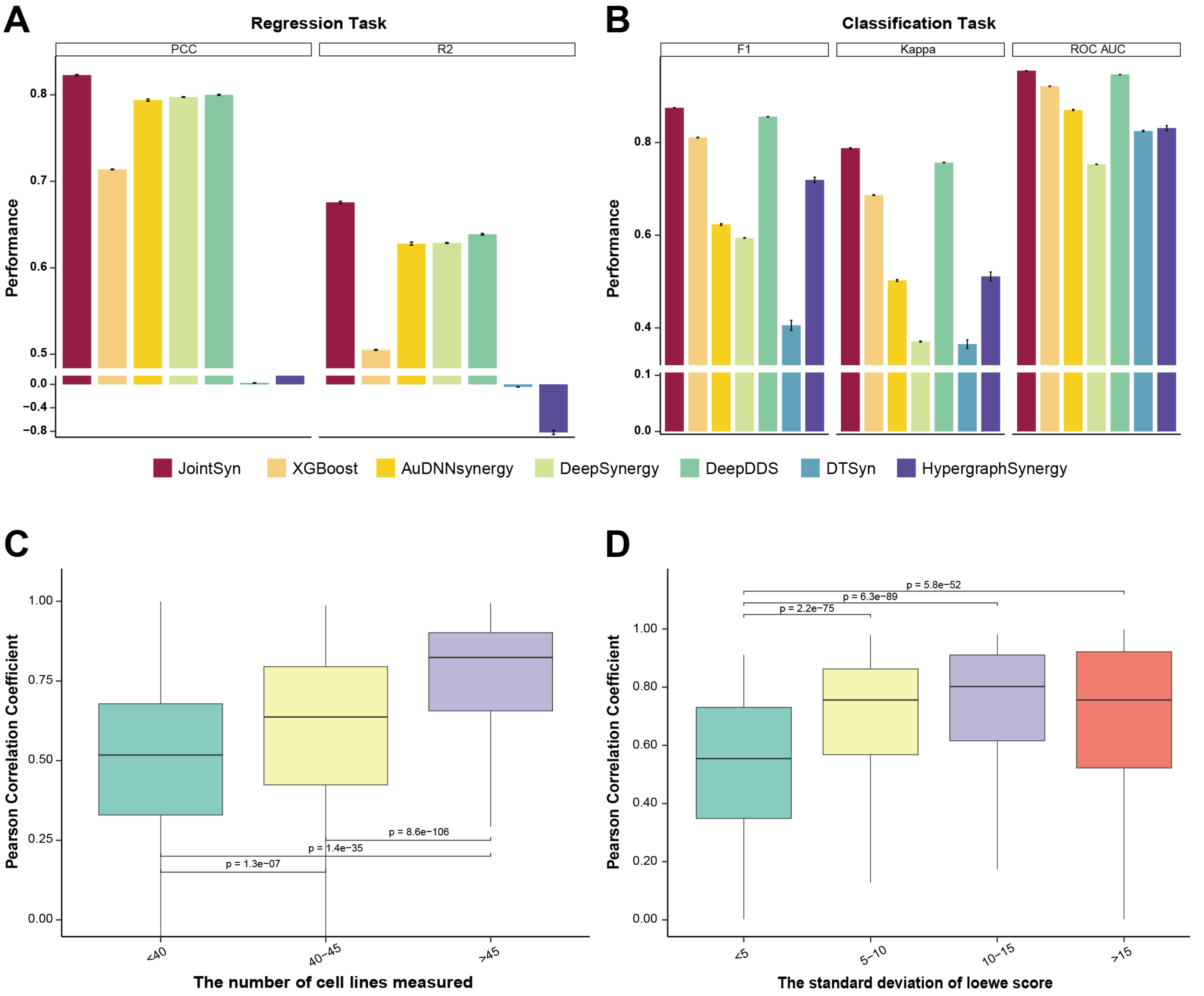


**Supplementary Figure 1 Evaluation of JointSyn on the NCI-ALMANAC benchmark dataset. a-b,** Performance comparison of JointSyn and other methods for the regression and classification tasks. Five-fold cross-validations were replicated 10 times to calculate the standard deviations (error bars). **c-d**, Taken PCC of O'Neil regression model as an example to discuss factors associated with the performance of each drug combination. **c**, The relationship between the PCC and the number of cell lines measured for each drug combination. **d**, The relationship between the PCC and the standard deviation of the real synergy scores.

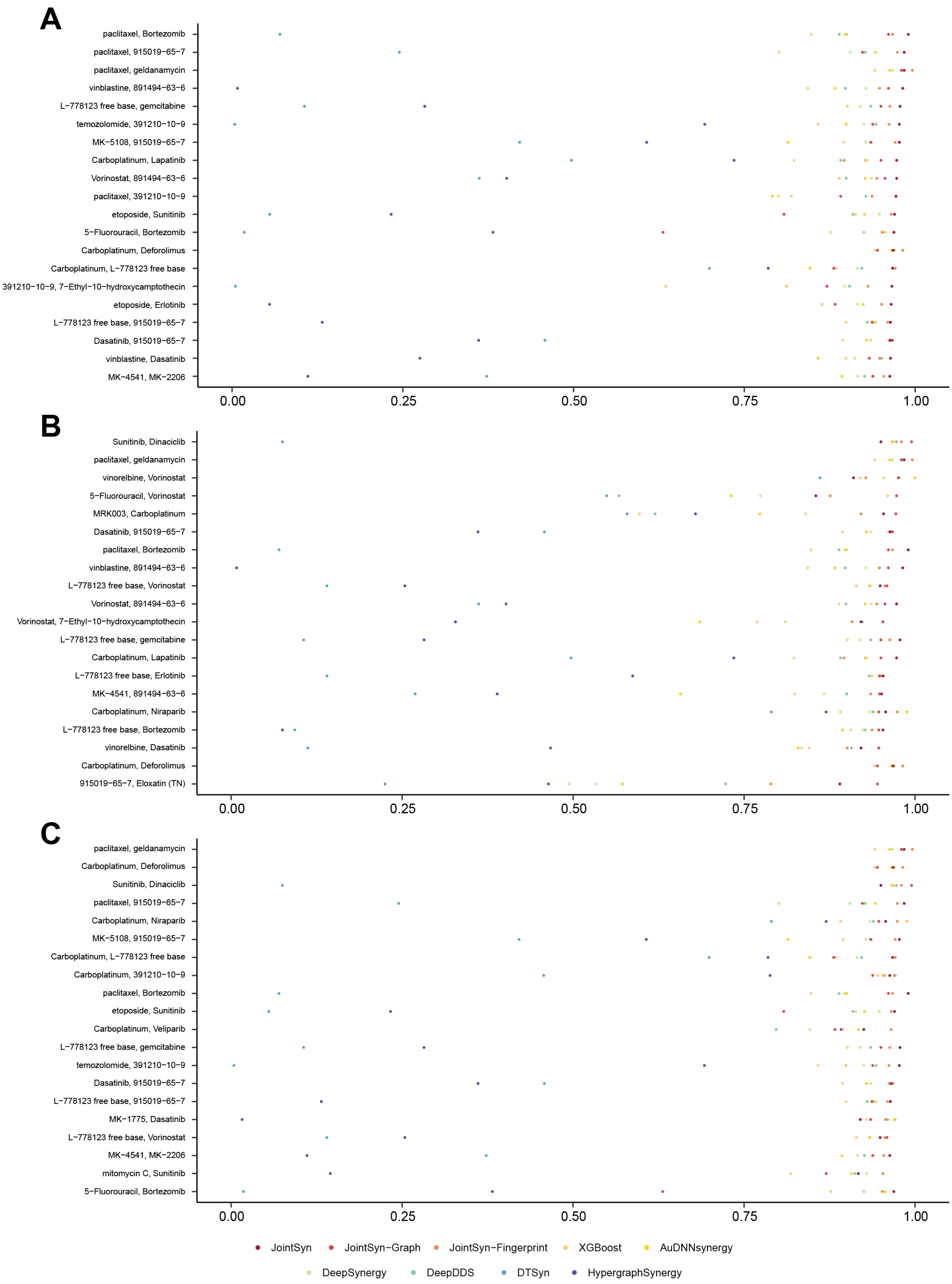


**Supplementary Figure 2 The contribution of two views by comparing them on each drug combination. a-c,** The top 20 drug combinations by the JointSyn, JointSyn- Graph and JointSyn-Fingerprint respectively.

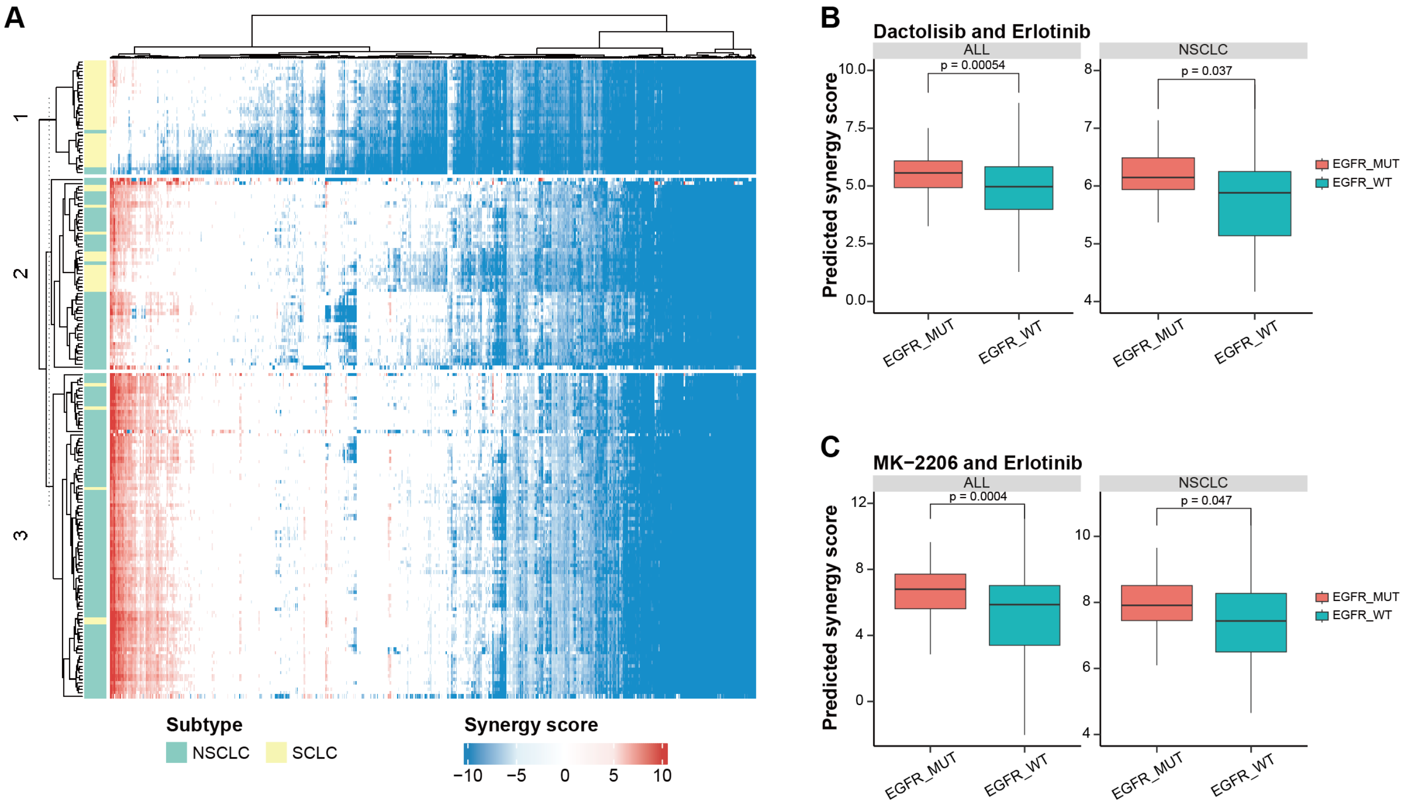


**Supplementary Figure 3 The predicted synergy scores of pan-cancers. a**, The predicted synergy scores of all drug combinations on lung cell lines. **b**, Comparison of the predicted synergy scores of dactolisib and erlotinib in cell lines with or without EGFR mutations. Pan-cancers and NSCLC were compared respectively. P-value was calculated by the Wilcoxon test. **c**, Comparison of the predicted synergy scores of MK-2206 and erlotinib in cell lines with or without EGFR mutations.
